# Effect of EMS induced mutation in rice cultivar Nagina 22 on salinity tolerance

**DOI:** 10.1101/2021.08.03.455004

**Authors:** Arun Shankar, OP Choudhary, Dharminder Bhatia, Kuldeep-Singh

## Abstract

Salinized hydroponic culture experiment with three salinity levels (EC control, 6 and 12 dS/m) was performed to screen salt tolerant mutants of aerobic rice cultivar Nagina 22 and to study the nature of salt tolerance from a total of 432 EMS induced M4 mutants. Plants were harvested 30 days after sowing. Growth parameters viz. root weight, shoot weight, root length, shoot length, Na and K concentrations in shoot and roots were measured. Combined Factor scores of growth parameters was computed by Principle Component Analysis using Minitab software.

At EC 12 dS/m 10 mutants out of 432 were able to survive. At moderate salinity, some mutant lines produced higher shoot weight compared to their respective control showing inverse trend and the effectiveness of EMS induced mutation in inducing salinity tolerance to these mutants. At high salinity only10 mutants survived (remained green) up to the time of 30 days harvest. These mutants performed well in terms of overall growth recording 2.1-2.5 times higher factor score and 8-14 times higher shoot weight compared to the N 22 check. One mutant N22-L-1010 almost completely excluded Na at xylem parenchyma level. Two other mutants N22-L-1013 and N22-L-806 maintained Na exclusion compared to the N22 check. N22 check and mutant N22-L-1009 maintained similar degree of Na exclusion though the N22 check died because it cannot maintain adequate K in shoot.

We conclude that EMS has induced salinity tolerance in some mutants. The study can be advanced further to characterize the putative mutants through molecular genetics approaches.

## Introduction

Soil salinity is a major abiotic threat limiting plant growth and development causing yield loss in many crops (Hasegawa et al., 2000; Qadir et al., 2007). Salt-affected soils contain excessive levels of water-soluble salts, especially sodium chloride (NaCl) (Tanji, 2002). Sodium chloride upon ionization produces sodium (Na^+^) and chloride (Cl^-^) ions causing ionic and osmotic stress in higher plants.(Mansour and Salama, 2000; Chinnusamy et al., 2005). Soil salinity is a very severe problem particularly in semiarid, and coastal rice-producing areas of the tropics (Lee et al., 2003). The rice production in the South Western districts of the state of Punjab, India which lies in semiarid zone is severely affected by soil salinity. The prime cause of salinity in this region is the brackish nature of underground water. Moreover, salinity is usually coupled with water logging due to shallow depth of water table. Hence the farmers prefer to grow rice as compared to other crops it can withstand standing water.

Considered moderately sensitive to salinity (Akbar et al., 1972; Mass and Hoffman, 1977; Mori et al., 1987) rice (*Oryza sativa* L. spp. *indica*) is one of the five main carbohydrate crops responsible for feeding the world’s population, and is especially important in Asian countries. Rice has previously been reported as being salt susceptible in both the seedling and reproductive stages (Zeng et al., 2001; Moradi and Ismail, 2007), leading to a reduction in yield of more than 50% when exposed to 6.65 dS m^-1^ ECe (Zeng and Shannon, 2000). Change in farming systems to alleviate salinity may be one crucial approach, however, it is likely to be a long and difficult process because it will require the use of new land and will not address the problem of growing crops in land that is already compromised (Yamaguchi and Blumwald 2005). Therefore, it is crucial to improve tolerance to salinity in rice by breeding high yielding salinity tolerant varieties.

Differential salt tolerance exist among different genera and species, and also within the different parts of the same species (Flowers and Hajibagheri 2001; Ismail, 2003). Studying the response of cultivars of any species to salinity provides a convenient and useful tool for understanding the basic mechanisms involved in salt tolerance (Tammam et al., 2008). Breeding rice varieties having in-built salt tolerance is realized as valuable, resource conserving, more economic and socially acceptable approach. Salt tolerance is a multigenic trait enabling plants to grow and sustain economic yield even when the salt concentration is high particularly NaCl (Hurkman, 1992; Deepa Sankar et al., 2011). Flowers and Yeo (1995) reported five ways to develop salt-tolerant crops: (1) developing halophytes as alternative crops; (2) use of interspecific hybridization to enhance the tolerance of current crops; (3) utilize the already present variation with respect to salt tolerance in existing crops; (4) develop variation within crops by using recurrent selection, mutagenesis or tissue culture, and (5) breeding for economic yield rather than tolerance. Conventional breeding of cultivars for salt tolerance is time consuming requiring multiple crosses for many generations and labor intensive (Peleman and Voort, 2003; Hwa and Yang, 2008). Moreover, traditional breeding methods are not suitable for screening a large sample size for exploiting variations in tolerance.

Chemical mutagens viz. EMS, DEB and sodium azide; irradiation viz. Gamma rays, X rays and fast neutrons have been used to generate large number of functional variations in rice. (Talebi et al, 2012). Chemicals generate mostly point mutations so they are ideal for producing missense and nonsense mutations providing a sequence of functional mutations. Hence, induced mutation technique to create variability among rice germplasm (Mba et al, 2007) and screening of salt resistant cultivars under solution culture (Gregorio et al., 1997) could become a viable alternative towards tolerance to soil salinity as screening of large number of germplasm under field conditions is difficult due to stress heterogeneity, presence of other soil-related stresses, and the significant influence of environmental factors (Gregorio et al., 1997). Most of the studies focus on the effects of sodium on the plant growth and the importance of control mechanisms for sodium uptake in salt-resistant plants, but little emphasis has been put on root selectivity for potassium viz a viz sodium (Asch et al 2000) which is discussed in this paper. Moreover, large numbers of papers has been published regarding salinity tolerance mechanisms yet very few salt tolerant cultivars has been released offering only slight improvement over parent lines (Flowers and Yeo, 1995; Glenn et al., 1999).

Keeping the above factors in mind, mutation was induced by EMS (Ethyl methane sulfonate) in aerobic rice cultivar Nagina 22 (which is also drought tolerant cultivar). A solution culture experiment was performed to screen salt tolerant mutants by studying their morphological parameters such as root, shoot length and dry weight and to study the underlying mechanism of salt tolerance by studying Na and K concentrations in root and shoot of the promising and worse mutant. Knowledge obtained from such studies will be helpful to characterize the putative mutants through molecular genetics approaches.

## Material and Methods

Salinized hydroponics culture experiment with three salinity levels (EC control, 6 and 12 dS/m) was conducted to screen salt tolerant mutant from a total of 432 EMS induced M_4_ mutants of drought tolerant rice cultivar Nagina 22. Plants were harvested 30 days after sowing. Root weight, shoot weight, root length, shoot length, Na and K concentrations in shoot and roots were measured. The experiment was conducted at the Greenhouse of School of Agricultural Biotechnology, Punjab Agricultural University Ludhiana situated at 300 56/ N latitude and 750 52/ E longitude and at an altitude of 247 m above mean sea level. Mutations in ‘Nagina 22’ were induced using 0.5% Ethyl methanesulfonate (EMS). These mutants were produced at IARI, New Delhi and distributed to all the collaborating centres in the country including Punjab Agricultural University. Nagina 22 (N22) was used as check.

### Creation of salinity and experimental set up

In control treatment no salts were added. Salts of NaCl and CaCl_2_.2H_2_O were added in 2:1 on equivalent basis to get the desired EC level. The electrical conductivity of the solution was monitored by the use of conductivity meter. Plastic trays (holding 8 litres of half Hoagland’s solution up to the brim) of the size 45 cm x 32 cm x 8 cm were used to hold nutrient solution. A thermocol sheet of the size 45 cm x 32 cm was cut and small holes of 2 cm diameter were made at distance of 5.5 cm (fig. 1). A plastic net was fixed on the lower side of the thermocol sheet to hold the seed in the respective holes which remained in touch with the Hoagland’s solution in the trays. Five seeds were placed in each hole. The Hoagland’s solution was replaced after 3 days. Electrical conductivity was checked periodically during the growth phase and maintained at targeted level. The composition of Hoagland solution is mentioned in Table 5.

**Fig. 1.**
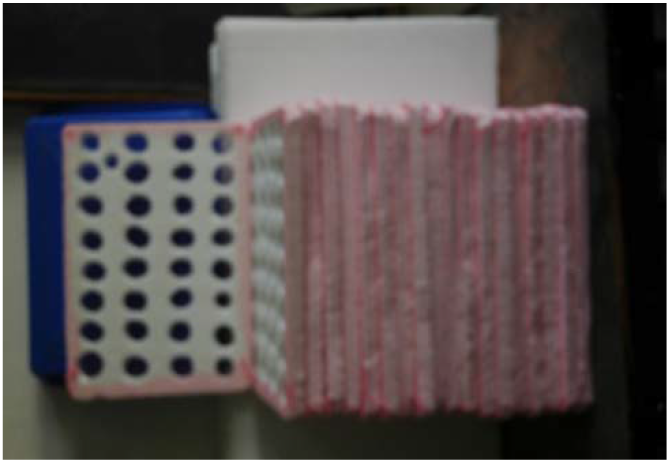
A Set of Sheets ready for Hydroponics trial.

**Fig. 2.**
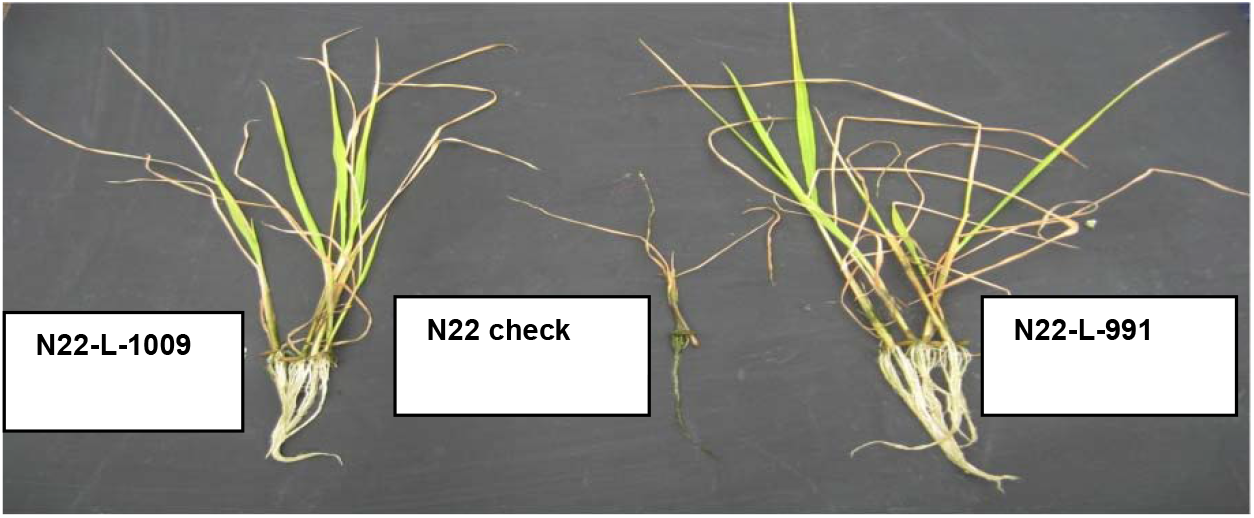
Tolerant mutants compared with N22 check at high salinity.

**Fig. 3.**
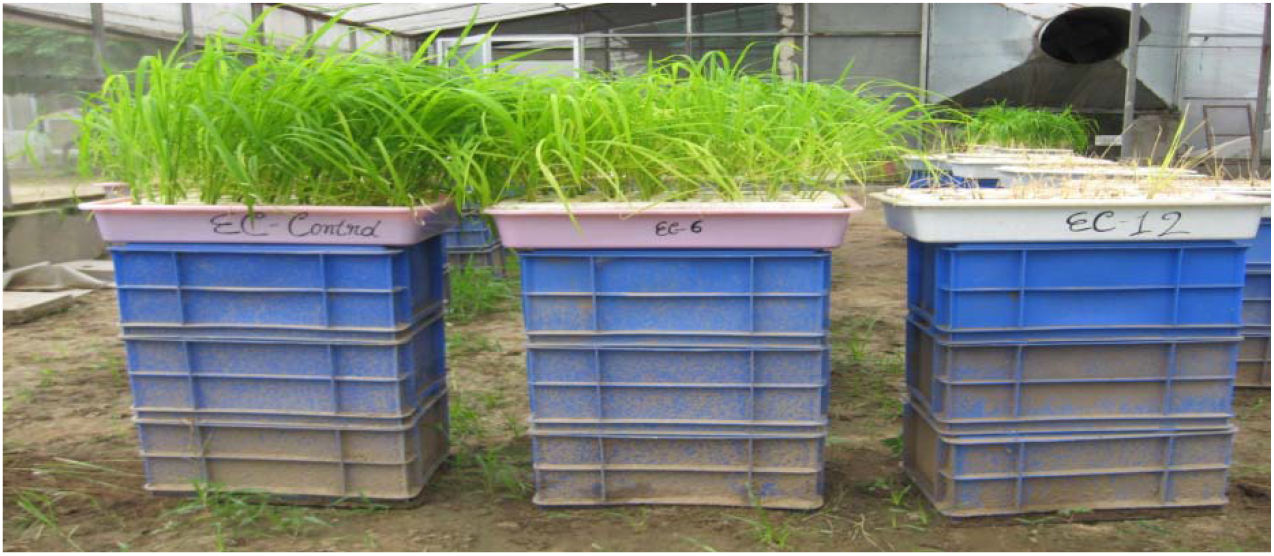
N22 mutants at three salinity levels

**Fig. 4.**
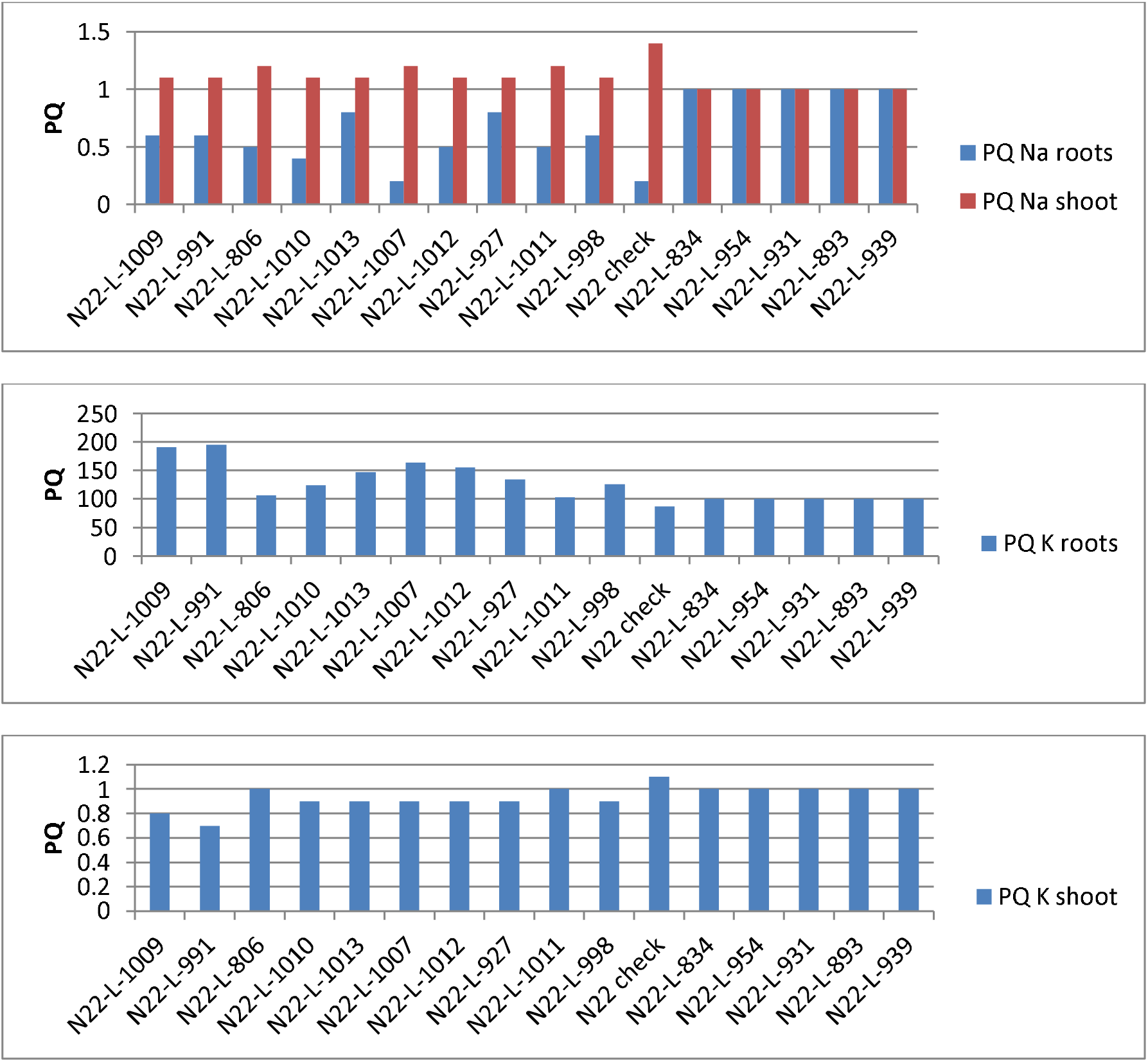
Partition quotient (PQ) of Na and K of some mutants in roots shoot at control EC

**Fig. 5.**
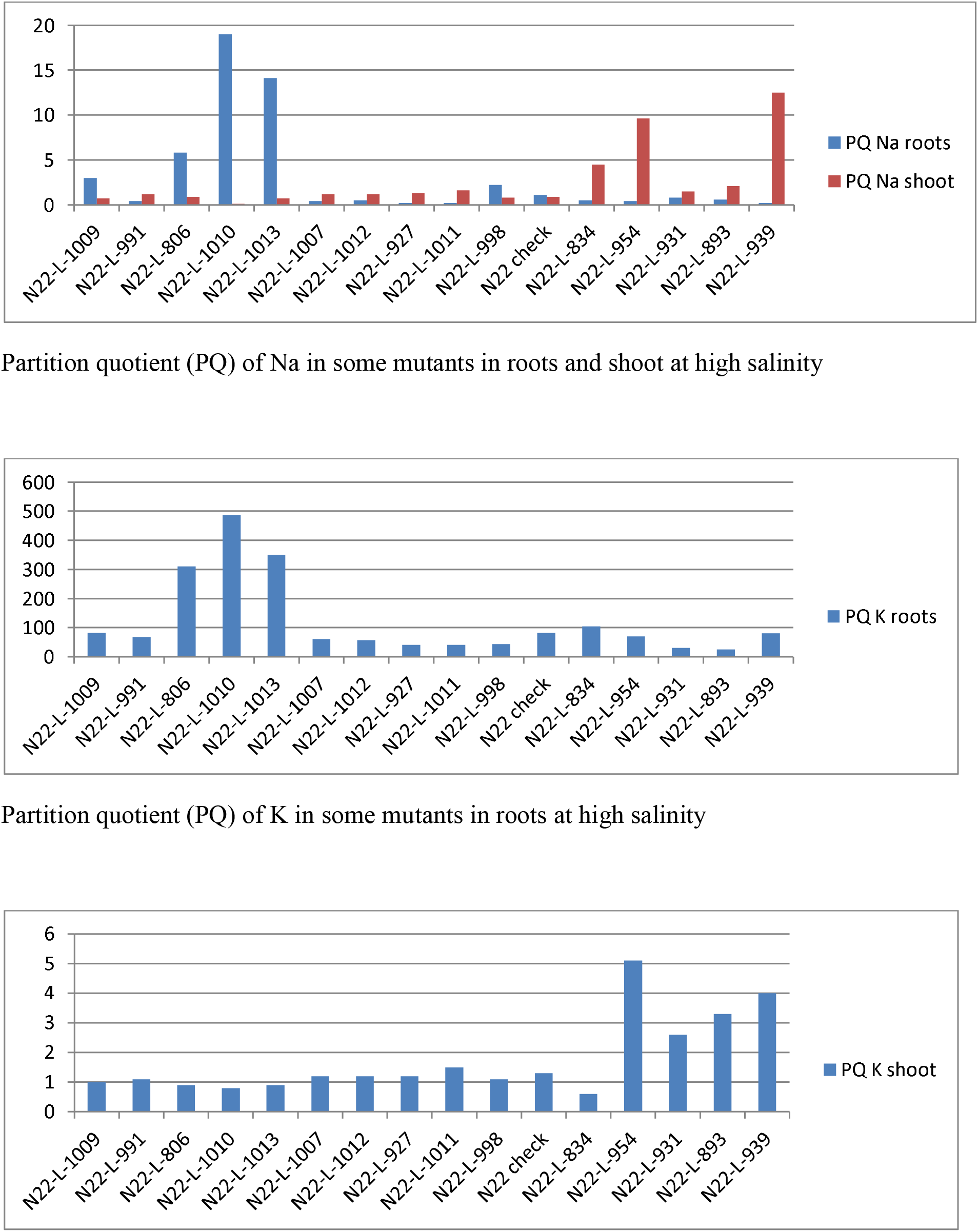
Partition quotient (PQ) of K in some mutants in shoot at high salinity

### Harvesting, collection and processing of shoot samples

Shoot and root length was recorded after 30 days of sowing and plants were harvested. The harvested shoot samples were washed with acidified deionized water, and then with distilled water and finally with double distilled water. The washed shoot samples were air dried and then dried in oven at 65±2°C. Thereafter, the shoots were weighed for dry matter yield and stored in paper bags. They were digested in diacid mixture of HNO_3_ and HClO_4_ in the ratio 3:1. The digested samples were analyzed for Na and K by flame photometer.

### Partition quotient

The dry weight of root and shoot was calculated as a percentage of total plant weight and mineral uptake of each organ was calculated as a percentage of total plant mineral uptake. Using these values, the normalized partitioning of that mineral within the plant was calculated by dividing each organ’s percentage mineral uptake by its percentage DW which will be denoted as partition quotient (PQ) (Waters and Grusak, 2008).

### Statistical analysis

Combined Factor scores of growth parameters was computed by Principle Component Analysis using Minitab software. The correlation analysis was done using data analysis tool in MS excel.

## Results and Discussion

The overall growth of screened mutants was quite good at control and EC 6 dS/m but the survival of most of the mutants was poor at EC 12 dS/m where only 10 mutants out of 432 were able to survive. According to Table 1, mutants were categorized based on shoot weight and factor score as good at moderate salinity (Sl. No. 1-10 and 17-24), good at high salinity (Sl. No. 1-10), worse at moderate salinity (Sl. No. 25-28) and worse at high salinity (Sl. No. 12-16). The N22 check was susceptible at moderate and high salinity. The combined factor score based on root length, root weight, shoot length and shoot weight (Table 1) was a good indicator of overall growth of mutants at different salinity levels. The combined factor score declined with salinity in all the mutants including N 22 check. This indicates that salinity generally imposes reduction to overall growth of plants. At moderate salinity, mutant lines (Sl. 17 to 24) produced higher shoot weight compared to their respective controls thereby showing inverse trend. These mutants performed well in terms of shoot length, root length and root weight (data not shown) as these mutants recorded 91-96 % relative factor score and more than twice shoot weight compared to their respective controls. Mutant lines N22-L-1240, N22-L-1195, N22-L-1247 and N22-L-1151 were the worse mutants at moderate salinity in terms of absolute shoot weight and factor score. N22 check was also susceptible compared to the tolerant mutants at moderate salinity performing better only compared to some worst mutants. These results are in accordance with Danaguiraman et al. (2003) who reported that, tolerant genotypes of rice recorded a higher germination percentage, root length, shoot length and vigour index.

**Table 1:** Growth parameters and factor score of N22 mutants at three salinity levels

| Sl. No. | Mutant ID | Low Salinity (EC control) |  |  | Moderate salinity (EC 6 dS m<sup>-1</sup>) |  |  | High salinity (EC 12 dS m<sup>-1</sup>) |  |  |
| --- | --- | --- | --- | --- | --- | --- | --- | --- | --- | --- |
|  |  | Shoot weight (mg) | Root weight (mg) | Factor Score | Shoot weight (mg) | Root weight (mg) | Factor Score | Shoot weight (mg) | Root weight (mg) | Factor Score |
| 1 | N22-L-1009 | 1022 | 269 | 97.1 | 702 | 209 | 79.3 | 212 | 37 | 40.5 |
| 2 | N22-L-991 | 1641 | 506 | 94.2 | 1013 | 236 | 76.4 | 220 | 75 | 41.1 |
| 3 | N22-L-806 | 913 | 340 | 95.8 | 818 | 135 | 76.2 | 208 | 5 | 36.0 |
| 4 | N22-L-1010 | 1230 | 298 | 94.8 | 846 | 139 | 81.1 | 318 | 15 | 40.5 |
| 5 | N22-L-1013 | 1195 | 298 | 94.0 | 861 | 294 | 81.1 | 185 | 4 | 36.9 |
| 6 | N22-L-1007 | 1200 | 262 | 91.8 | 863 | 232 | 74.2 | 185 | 75 | 41.1 |
| 7 | N22-L-1012 | 1181 | 235 | 91.3 | 968 | 220 | 78.8 | 211 | 75 | 34.2 |
| 8 | N22-L-927 | 1454 | 510 | 83.4 | 970 | 127 | 66.1 | 145 | 51 | 36.5 |
| 9 | N22-L-1011 | 827 | 262 | 91.8 | 1079 | 230 | 77.5 | 190 | 150 | 35.8 |
| 10 | N22-L-998 | 907 | 296 | 95.2 | 1297 | 283 | 80.2 | 248 | 53 | 37.5 |
| 11 | **N22 check** | 512 | 266 | 53.9 | 135 | 61 | 39.7 | 18 | 27 | 16.3 |
| 12 | N22-L-834 | 1022 | 133 | 73.2 | 273 | 79 | 57.4 | 4 | 26 | 19.1 |
| 13 | N22-L-954 | 906 | 297 | 78.1 | 360 | 274 | 63.1 | 2 | 27 | 19.7 |
| 14 | N22-L-931 | 1429 | 388 | 83.7 | 563 | 211 | 66.9 | 6 | 14 | 21.8 |
| 15 | N22-L-893 | 752 | 231 | 80.6 | 345 | 206 | 63.9 | 7 | 21 | 23.7 |
| 16 | N22-L-939 | 1123 | 343 | 86.2 | 385 | 186 | 71.7 | 3 | 46 | 21.9 |
| 17 | N22-L-1109 | 356 |  | 64.9 | 930 |  | 62.0 |  |  |  |
| 18 | N22-L-1111 | 293 |  | 61.0 | 918 |  | 57.0 |  |  |  |
| 19 | N22-L-1094 | 306 |  | 62.5 | 886 |  | 59.5 |  |  |  |
| 20 | N22-L-1098 | 394 |  | 68.5 | 825 |  | 64.2 |  |  |  |
| 21 | N22-L-1092 | 391 |  | 67.0 | 809 |  | 61.9 |  |  |  |
| 23 | N22-L-1108 | 295 |  | 69.1 | 751 |  | 63.0 |  |  |  |
| 24 | N22-L-1090 | 316 |  | 63.6 | 750 |  | 58.9 |  |  |  |
| 25 | N22-L-1240 | 313 |  | 52.2 | 59 |  | 35.7 |  |  |  |
| 26 | N22-L-1195 | 125 |  | 46.0 | 52 |  | 36.0 |  |  |  |
| 27 | N22-L-1247 | 396 |  | 50.7 | 49 |  | 36.6 |  |  |  |
| 28 | N22-L-1151 | 438 |  | 52.0 | 47 |  | 36.8 |  |  |  |

According to Table 2, K uptake gradually decreased with increase in salinity level except for the mutants showing inverse trend at moderate salinity (Sr. No. 17-24) while shoot Na uptake generally increased with salinity in all the mutants. Shoot Na:K ratio increased in all the mutants compared to their respective controls. However, increase was the maximum in the succeptible mutants (Sr. No.25-28) showing 1.2-2 times increase compared to the N22 check. Conversely, the good mutants (Sr. No. 1-10 and 17-24) recorded 30-76% lower Na:K ratio compared to N22 check. Shoot weight was highly negatively correlated with shoot Na:K ratio at moderate and high salinity (r^2^ = −0.82 and r^2^ = −0.74, respectively). The relationship between potassium and sodium uptake by the rice plant and tolerance to salinity stress has been reported by Qadar, (1988); Pandey & Srivastava, (1991); Bohra & Dörffling, (1993).

**Table 2:** Shoot Na and K uptake of some mutants at low, moderate and high salinity levels

| Sl. No. | Mutant ID | Low Salinity (EC control) |  |  |  | Moderate salinity (EC 6 dS m<sup>-1</sup>) |  | High salinity (EC 12 dS m<sup>-1</sup>) |  |  |  |
| --- | --- | --- | --- | --- | --- | --- | --- | --- | --- | --- | --- |
|  |  | Shoot Na uptake | Shoot K uptake | Root Na uptake | Root K uptake | Shoot Na uptake | Shoot K uptake | Shoot Na uptake | Shoot K uptake | Root Na uptake | Root K uptake |
| 1 | N22-L-1009 | 11.2 | 275 | 1.6 | 182 | 73.7 | 214 | 35.4 | 86 | 28.3 | 11.9 |
| 2 | N22-L-991 | 23.3 | 315 | 3.8 | 268 | 187 | 330 | 530 | 62 | 63.6 | 12.8 |
| 3 | N22-L-806 | 14.7 | 379 | 2.2 | 154 | 112 | 249 | 337 | 163 | 52.7 | 12.8 |
| 4 | N22-L-1010 | 18.5 | 451 | 1.6 | 143 | 87.1 | 308 | 59.8 | 167 | 361 | 46.8 |
| 5 | N22-L-1013 | 15.3 | 424 | 2.7 | 176 | 68.0 | 316 | 353 | 104 | 150 | 8.3 |
| 6 | N22-L-1007 | 23.4 | 308 | 1.1 | 128 | 138 | 283 | 596 | 54.6 | 84.8 | 11.6 |
| 7 | N22-L-1012 | 18.0 | 417 | 1.6 | 145 | 147 | 315 | 333 | 86.5 | 55.4 | 15.1 |
| 8 | N22-L-927 | 20.1 | 429 | 5.4 | 228 | 93.1 | 342 | 529 | 94.8 | 22.8 | 11.5 |
| 9 | N22-L-1011 | 17.9 | 455 | 2.2 | 150 | 69.1 | 323 | 304 | 82.7 | 36.9 | 18.3 |
| 10 | N22-L-998 | 13.6 | 331 | 2.2 | 149 | 162 | 409 | 87 | 126 | 53.3 | 10.6 |
| 11 | **N22 check** | 13.3 | 428 | 1.1 | 181 | 81 | 99.3 | 22.3 | 3.9 | 42.9 | 3.8 |
| 12 | N22-L-834 | 36.8 | 413 | 4.8 | 54 | 41.2 | 48.0 | 60.3 | 1.6 | 40.8 | 17.0 |
| 13 | N22-L-954 | 15.2 | 418 | 5.0 | 137 | 69.1 | 90.0 | 69.0 | 3.3 | 35.3 | 6.1 |
| 14 | N22-L-931 | 54.3 | 1292 | 14.7 | 351 | 79.9 | 124 | 52.7 | 4.8 | 65.2 | 1.3 |
| 15 | N22-L-893 | 20.1 | 382 | 6.2 | 117 | 42.8 | 70.0 | 55.4 | 4.2 | 48.9 | 1.0 |
| 16 | N22-L-939 | 10.9 | 356 | 3.3 | 109 | 45.8 | 90.1 | 66.3 | 2.9 | 20.1 | 9.0 |
| 17 | N22-L-1109 | 20.6 | 390 | - | - | 259 | 789 | - | - | - | - |
| 18 | N22-L-1111 | 10.3 | 200 | - | - | 360 | 666 | - | - | - | - |
| 19 | N22-L-1094 | 14.1 | 370 | - | - | 282 | 764 | - | - | - | - |
| 20 | N22-L-1098 | 15.2 | 308 | - | - | 281 | 870 | - | - | - | - |
| 21 | N22-L-1092 | 15.2 | 417 | - | - | 73.6 | 377 | - | - | - | - |
| 23 | N22-L-1108 | 17.4 | 253 | - | - | 238 | 701 | - | - | - | - |
| 24 | N22-L-1090 | 4.4 | 62.3 | - | - | 237 | 633 | - | - | - | - |
| 25 | N22-L-1240 | 7.5 | 280 | - | - | 140 | 87 | - | - | - | - |
| 26 | N22-L-1195 | 7.1 | 140 | - | - | 87 | 84 | - | - | - | - |
| 27 | N22-L-1247 | 13.5 | 428 | - | - | 103 | 102 | - | - | - | - |
| 28 | N22-L-1151 | 7.0 | 99 | - | - | 101 | 96 | - | - | - | - |

Plants use three main mechanisms to survive salinity stress: osmotic tolerance, Na^+^ exclusion and Na^+^ tissue tolerance (Munns and Tester 2008). Earlier Cl^-^ was thought to be main cause of salt injuries but later it was confirmed that excessive uptake of Na^+^ is the chief cause of salt injuries to plants (Clarkson and Hanson 1980). Salt injuries include injuries related to osmotic effects (especially in the initial growth stages (Lauchli and Grattan, 2007). Hence, exclusion of the sodium ion (Na^+^) from shoot and increased absorption of K^+^ to maintain a good Na^+^: K^+^ balance in the shoot are very important traits of tolerant genotypes (Roshandel and Flowers 2009).

Blumwald (2000) reported that ion ratios in plants are altered by the influx of Na^+^ through K^+^ pathways. The similarity of the hydrated ionic radii of Na^+^ and K^+^ makes it difficult to discriminate between them and this is the basis of Na^+^ toxicity. However, at moderate salinity, most of the mutants marinated fairly lower Na:K ratio (Table 2) except for the sensitive mutants (Sr. 25-28) leading to their better performance compared to the sensitive mutants. Kader and Lindberg (2008) reported that the transient uptake of Na^+^, which occurs only in the tolerant cultivar (Na exclusion), and the fast compartmentalization of Na^+^ into the vacuole (tissue tolerance), probably are the most important cellular trait for Na^+^ tolerance in rice. Later, Munns and Tester (2008) in their review article suggested that Na exclusion in shoots is associated with partitioning of Na to roots while tissue tolerance is associated with compartmentalization of Na^+^ into the vacuole and Na transport can be controlled by tolerant plants at moderate salinity. Hence, the susceptibility of worse mutants to moderate salinity could be explained largely by their higher Na:K ratio compared to the N22 check. Roshandel and Flowers (2009) suggested that salt-tolerant cultivars of rice accumulate less Na^+^ and more K^+^ in comparison to susceptible ones. Results of this study reveal that at moderate salinity, plants adopt sodium avoidance mechanism by excluding sodium to maintain a low Na:K ratio. Also Munns (2002) has suggested that Na to K selectivity of plants roots minimizing the entry of Na into plants and maintaining effective uptake of K together to the shoots plays crucial role in salinity tolerance to plants.

At high salinity, only 10 mutants survived (remained green) up to the time of 30 days harvest (Sr. No. 1-10). The N 22 check did not survive. These mutants performed well in terms of overall growth recording 2.1-2.5 times higher factor score and 8-14 times higher shoot weight compared to the N 22 check. Five of the worse mutants (Sr. No. 12-16, Table 1) at high salinity recorded 61-89 % lower shoot weight compared to N22 check. At high salinity, the shoot Na:K ratio ranged between 0.36 to 10.92 for the good mutants (Sr. No. 1-10, Table 2) and 10.98-37.69 for the worst mutants (Sr. No. 12-16, Table 2). The N22 check recorded a value of 5.72. The other worse mutants died because they cannot maintain Na exclusion at the xylem parenchyma level. One mutant N22-L-1010 almost completely excluded Na at xylem parenchyma level. Two other mutants N22-L-1013 and N22-L-806 maintained Na exclusion compared to the N22 check. N22 check and mutant N22-L-1009 maintained similar degree of Na exclusion though the N22 check died because it cannot maintain adequate K in shoot from roots at the xylem parenchyma level. When salinity results from an excess of NaCl, the most common type of salt stress, the concentrations of sodium (Na^+^) and chloride (Cl^-^) in the plant increase, and concentrations of potassium (K^+^) and calcium (Ca^+^) are reduced. This is known as salt-specific or ion-excess effect of salinity. Increases in salt concentrations more than optimum value can lead to toxic accumulation of ions such as Cl^-^ and, in particular, Na^+^ exerting ion toxicity at cellular level with varying concentrations according to the capacity of salt tolerance of plants (Diedhiou, 2006).

Exclusion of Na from leaves is facilitated by two mechanisms: first, an outward active transport of Na at the plasma membrane of root cortical cells resulting into extrusion of Na to the outward medium (Jeschke, 1984, Pitman and Saddler, 1967). Second, Resorption of Na ions from the xylem sap and its accumulation by xylem parenchyma cells in the roots and stem base reducing amount of Na reaching the leaves (Lauchli, 1984). In this study, three mutants N22-L-1010, N22-L-1009 and N22-L-998 maintained lesser relative Na: K Partition Quotient (PQ) ratio compared to N22 check indicating Na to K discrimination at the xylem parenchyma level. Notably, most of the other good mutants survived in spite of having higher relative Na: K PQ ratio than N22 check. This is in accordance to the Na tissue tolerance mechanism. All the worse mutants maintained higher relative Na: K PQ ratio compared to N22 check showing poor Na: K discrimination. Sodium accumulation at toxic levels results into necrosis of leaves, shortening of lifespan of leaves, growth and yield reductions associated to the capacity of Na to compete with K for binding sites essential for cellular function (Tester and Davenport, 2003).

According to Table 3, PQ of K in roots at high salinity decreased for six mutants and N22 check compared to their respective controls while four mutants N22-L-806, N22-L-1010, N22-L-1013 and N22-L-834 recorded an increase. The N22 check recorded lowest PQ of K in shoots while all the good mutants recorded 1.5-2.5 times higher PQ of K in shoots compared to the N 22 check. Notably, mutants characterized as worse recorded higher (4.3-8.5 times) PQ of K in shoots compared to N22 check. Under control EC, the N22 check recorded higher PQ of Na in shoots compared to all other mutants (good or worse). Regarding PQ of Na in shoots, all the mutants accumulated either higher or similar Na compared to N22 check except one mutant line, N22-L-1010 which recorded 90% lower shoot Na accumulation compared to N22 check. However, the worse mutants on the other hand recorded 2.3 to 13.9 times higher shoot Na accumulation except one mutant N22-L-931(1.7 times) while the good mutants recorded (0.1-1.8 times). Among the good mutants, N22-L-1009, N22-L-1010, N22-L-1013 and N22-L-998 recorded 22, 89, 22 and 11 % lesser Na in shoots compared to N22 check. N22 check died at shoot Na PQ value of 0.9. Considering this value as the threshold value all the worse mutants accumulated higher shoot Na PQ leading to their poor performance at high salinity.

**Table 3:** Partition quotient of Na in shoot and root of some mutants at high salinity and their respective controls

| Sl. No. | Mutants | EC control |  |  |  | EC 12 (dS/m) |  |  |  |
| --- | --- | --- | --- | --- | --- | --- | --- | --- | --- |
|  |  | PQ Na<br>roots | PQ K<br>roots | PQ Na<br>shoot | PQ K<br>shoot | PQ Na<br>roots | PQ K<br>roots | PQ Na<br>shoot | PQ K<br>shoot |
| 1 | N22-L-1009 | 0.6 | 191 | 1.1 | 0.8 | 3.0 | 81.7 | 0.7 | 1.0 |
| 2 | N22-L-991 | 0.6 | 195 | 1.1 | 0.7 | 0.4 | 67.4 | 1.2 | 1.1 |
| 3 | N22-L-806 | 0.5 | 106 | 1.2 | 1.0 | 5.8 | 310 | 0.9 | 0.9 |
| 4 | N22-L-1010 | 0.4 | 124 | 1.1 | 0.9 | 19.0 | 486 | 0.1 | 0.8 |
| 5 | N22-L-1013 | 0.8 | 147 | 1.1 | 0.9 | 14.1 | 350 | 0.7 | 0.9 |
| 6 | N22-L-1007 | 0.2 | 164 | 1.2 | 0.9 | 0.4 | 60.5 | 1.2 | 1.2 |
| 7 | N22-L-1012 | 0.5 | 155 | 1.1 | 0.9 | 0.5 | 56.6 | 1.2 | 1.2 |
| 8 | N22-L-927 | 0.8 | 134 | 1.1 | 0.9 | 0.2 | 41.7 | 1.3 | 1.2 |
| 9 | N22-L-1011 | 0.5 | 103 | 1.2 | 1.0 | 0.2 | 41.1 | 1.6 | 1.5 |
| 10 | N22-L-998 | 0.6 | 126 | 1.1 | 0.9 | 2.2 | 44.1 | 0.8 | 1.1 |
| **11** | **N22 check** | 0.2 | 86.8 | 1.4 | 1.1 | 1.1 | 82.6 | 0.9 | 1.3 |
| 12 | N22-L-834 | 1.0 | 100.0 | 1.0 | 1.0 | 0.5 | 105 | 4.5 | 0.6 |
| 13 | N22-L-954 | 1.0 | 99.9 | 1.0 | 1.0 | 0.4 | 69.7 | 9.6 | 5.1 |
| 14 | N22-L-931 | 1.0 | 100.0 | 1.0 | 1.0 | 0.8 | 30.2 | 1.5 | 2.6 |
| 15 | N22-L-893 | 1.0 | 100.0 | 1.0 | 1.0 | 0.6 | 24.9 | 2.1 | 3.3 |
| 16 | N22-L-939 | 1.0 | 100.0 | 1.0 | 1.0 | 0.2 | 80.5 | 12.5 | 4.0 |

The PQ of Na in shoots was lower in good mutants compared to worse mutants. Intracellular K^+^/Na^+^ balance is an important factor in salt tolerance, because selectively taking up and maintaining a high level of K^+^ restricts the entry of Na^+^ (Blumwald 2000). According to Table 4, one mutant N22-L-1010 almost completely excluded Na at xylem parenchyma level through excluding Na from shoots. Two other mutants N22-L-1013 and N22-L-806 also maintained shoot Na exclusion compared to the N22 check. This mode of Na exclusion has been studied in *Triticeae* (Greenway and Munns, 1980; Gorham et al., 1985) and barley (Forster et al., 1994).

**Table 4:** Partitioning of Na and K in shoots compared to their respective controls

| Mutant lines | Relative Na PQ shoots: roots | Relative K PQ shoots: roots |
| --- | --- | --- |
| N22-L-1009 | 0.127 | 2.921 |
| N22-L-991 | 1.636 | 4.544 |
| N22-L-806 | 0.065 | 0.308 |
| N22-L-1010 | 0.002 | 0.226 |
| N22-L-1013 | 0.036 | 0.421 |
| N22-L-1007 | 0.500 | 3.619 |
| N22-L-1012 | 1.091 | 3.656 |
| N22-L-927 | 4.727 | 4.269 |
| N22-L-1011 | 3.333 | 3.755 |
| N22-L-998 | 0.198 | 3.498 |
| N22 check | 0.117 | 1.242 |
| N22-L-834 | 9.000 | 0.569 |
| N22-L-954 | 24.000 | 7.310 |
| N22-L-931 | 1.875 | 8.609 |
| N22-L-893 | 3.500 | 13.253 |
| N22-L-939 | 62.500 | 4.969 |

**Table 5:** Composition of Hoagland’s solution used:

|  |  |  |
| --- | --- | --- |
| The following Stock solutions were made: |  |  |
| 1. Ca (NO<sub>3</sub>)<sub>2</sub>.4H<sub>2</sub>O | - | 236.1 g/l |
| 2. KNO<sub>3</sub> | - | 101.1 g/l |
| 3. KH<sub>2</sub>PO<sub>4</sub> | - | 136.1 g/l |
| 4. MgSO<sub>4</sub>.7H<sub>2</sub>O | - | 246.5 g/l |
| 5. Trace elements (made up to 1 L) |  |  |
| H<sub>3</sub>BO<sub>3</sub> | - | 2.8 g |
| MnCl<sub>2</sub>.4H<sub>2</sub>O | - | 1.8 g |
| ZnSO<sub>4</sub>.7H<sub>2</sub>O | - | 0.2 g |
| CuSO<sub>4</sub>.5H<sub>2</sub>O | - | 0.1 g |
| NaMoO<sub>4</sub> |  | 0.025 g |
| 6. FeEDTA |  |  |
| EDTA.2Na | - | 10.4 g |
| FeSO<sub>4</sub>.7H<sub>2</sub>O | - | 7.8 g |
| KOH | - | 56.1 g |
| To make up 1 L of KOH, pH was adjusted to ~5.5 using H<sub>2</sub>SO<sub>4</sub>. Then EDTA.2Na and FeSO<sub>4</sub>.7H<sub>2</sub>O were added. |  |  |
| To make 1 L half Hoagland's solution from these stocks, following quantities were added |  |  |
| 7 ml Ca (NO<sub>3</sub>)<sub>2</sub> stock |  |  |
| 5 ml KNO<sub>3</sub> |  |  |
| 2ml KH<sub>2</sub>PO<sub>4</sub> |  |  |
| 2 ml MgSO<sub>4</sub> |  |  |
| 1 ml Trace elements |  |  |
| 1 ml FeEDTA |  |  |
| to 1 L water |  |  |

The characteristic of Na exclusion mechanism for salinity tolerance is lower Na accumulation in shoots by partioning excess Na to roots and this does not involve higher K uptake (Munns and Testor, 2008). As evident from Table 4, the mutants N22-L-1013, N22-L-806 and N22-L-1010 excluded Na in the shoots to survive. At the same time it was not accompanied by K accumulation in shoots indicating these mutants are true Na excluders. The worst mutants were not able to exclude Na from shoots hence they could not survive. The N22 check was not able to maintain adequate Na:K ratio and performed poorly.

## Conclusions

Based on our studies we conclude that EMS has induced variation in mutants with respect to their response to different levels of salinity and that can be exploited to identify the tolerant as well as susceptible mutants to salinity. N22-L-1013, N22-L-806 and N22-L-1010 were identified as typical Na excluders while rest of the tolerant mutants showed tissue tolerance mechanism.

## Acknowledgment

Authors are highly acknowledging the financial support provided by the Indian Council of Agricultural research (ICAR) in conducting this research.

